# Trio deep-sequencing does not reveal unexpected mutations in Cas9-edited monkeys

**DOI:** 10.1101/339143

**Authors:** Xin Luo, Yaoxi He, Chao Zhang, Xiechao He, Lanzhen Yan, Min Li, Ting Hu, Yan Hu, Jin Jiang, Xiaoyu Meng, Weizhi Ji, Xudong Zhao, Ping Zheng, Shuhua Xu, Bing Su

## Abstract

CRISPR-Cas9 is a widely-used genome editing tool, but its off-target effect remains a concern, especially in view of future clinical applications. Non-human primates (NHPs) share close genetic and physiological similarities with humans, making them an ideal preclinical model for developing Cas9-based therapies. However, no comprehensive *in vivo* off-target assessment has been conducted in NHPs. Here we performed whole genome trio sequencing of Cas9-treated monkeys. We found they only carried a small number of *de novo* mutations that can be explained by expected spontaneous mutations, and no unexpected mutations were detected.

---

CRISPR-Cas9 has been widely used to facilitate efficient genome editing in model and non-model animals^1^. The specificity of CRISPR-Cas9 relies on the designed 20bp guide RNA (sgRNA) and PAM^2, 3^. In the genome, there are often many sgRNA-like sequences. Consequently, CRISPR-Cas9 may generate nonspecific editing, leading to off-target effects.

CRISPR-Cas9 is a promising tool for correcting deleterious mutations causing human genetic diseases. However, the potential off-target effect makes it unsafe to implement in clinical therapeutic settings. Previously, most of the studies on off-target were carried out in rodents or human cells. Rodents serve as important animal models in pre-clinical studies. However, rodents have failed to show features of human disorders in many aspects. For example, the clinical symptoms involving high level cognitive functions cannot be reproduced faithfully in rodent models^4^. By contrast, non-human primates (NHPs) are genetically and physiologically similar with humans. Macaque monkeys have been used in biomedical research and are among the highest primates that can be genetically manipulated (without serious ethical concerns) to construct preclinical models for Cas9-based therapies^5,6^. Hence, exploring the off-target activity in Cas9-edited monkeys becomes crucial for future clinical applications.

The off-target effect has been investigated using whole genome sequencing (WGS) of Cas9-edited cells or animals^7–11^. However, previous studies mainly focused on the potential off-target loci predicted by sgRNA binding, not on a genome-wide evaluation of *de novo* mutations (DNMs). Recently, Schaefer et al. found plenty unexpected mutations using WGS of Cas9-edited mice^12^ though the claim was challenged by several groups^13–18^, and the latest trio sequencing of Cas9-edited mice did not see unexpected off-target activity^13^. To evaluate the situation in monkeys, we performed trio WGS of Cas9-edited rhesus monkeys (*Macaca mulatta*). We also analyzed the published trio WGS data of Cas9-edited cynomolgus monkeys (*Macaca fascicularis*)^19^.

We designed two sgRNAs to target exon2 and exon4 of *MCPH1*- a human autosomal recessive primary microcephaly gene^20^ (Supplementary Fig. 1a). Firstly, using zygotic injection of Cas9 mRNA and two sgRNAs, we achieved a high knockout efficiency for *MCPH1* at embryo level. Among the 15 rhesus monkey embryos tested, 13 of them were knockout positive (86.6%), including 3 (20%) knockout homozygotes (Supplementary Fig. 2, 3). To generate *MCPH1* knockout monkeys, we microinjected 30 zygotes, among which 24 zygotes developed normally and were transferred into 6 surrogate females, resulting in two pregnancies of twins and triplets, respectively. The surrogate female with twins had premature delivery at 138-days gestation, leading to a live male monkey and a dead female monkey. We performed C-section at 160-days gestation for the other surrogate female with triplets, and all three monkeys (one male and two females) were alive (Supplementary Fig. 1b, Supplementary Table 1). We used PCR-clone sequencing to evaluate the Cas9-editing status, and the result showed that all offspring monkeys were successfully modified by Cas9 except for rmO4^*ko*\*^ from the triplets. The dead female monkey (rmO2^*ko*^) was a homozygous knockout (Supplementary Fig. 1c). Interestingly, all Cas9-induced mutations were located in exon2 of *MCPH1.* No exon4 mutations were detected although sgRNA2 was designed to target exon4 and showed a high efficiency in the embryo test (Supplementary Fig. 3).

The five Cas9-treated monkeys and their three wildtype parents were subject to WGS by the Illumina X10 platform (Methods) (Fig. 1a, Supplementary Table. 1). Blood samples were taken from the four live monkeys for DNA extraction, and for the dead monkey, multiple tissues (brain, liver and muscle) were sampled to represent different germ layers. We achieved a median 46× depth of genome coverage (Table 1). The WGS data exhibited a high reads quality with Q30>85%, mean duplicate percentage of 12.03% and properly-paired reads >96% (Supplementary Table 2).

**Fig. 1.**
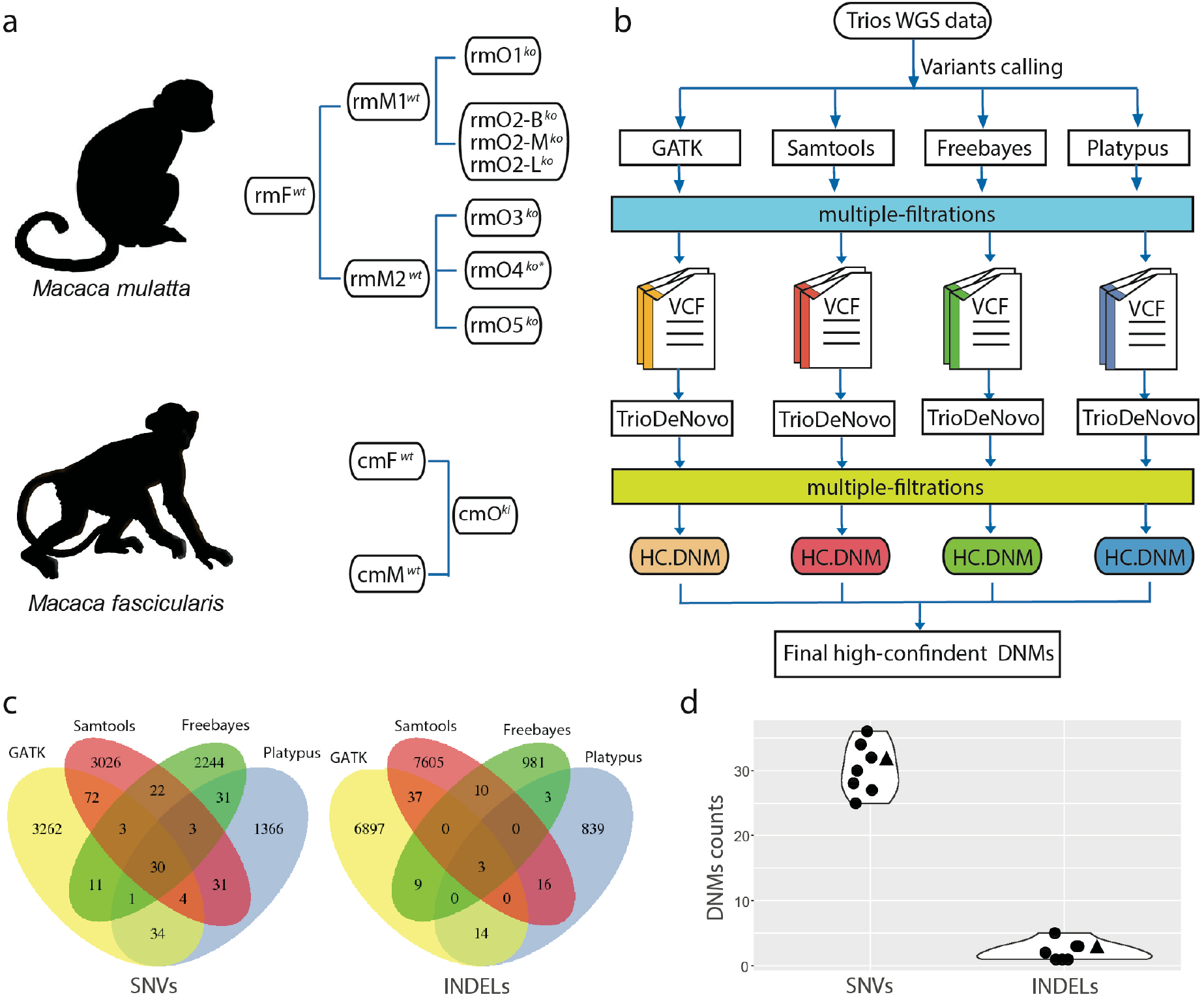
Detecting *de novo* mutations (DNMs) in the Cas9-treated monkeys. **a,** The analyzed trios of rhesus monkeys (rm) and cynomolgus monkey (cm). F-father, M-mother, O-offspring. ko-knockout, ki-knockin. B-brain, L-liver, and M-muscle; **b**, The pipeline of variant calling, filtering and DNMs identification. HC.DNM-high confident DNMs. **c,** Venn diagram of the high-confident DNMs for rmO2-B^*ko*^ identified by overlapping candidate DNMs from four different calling tools. **d,** Summary of the identified high-confident DNMs. The circles refer to Ca9-treated rhesus monkeys and the triangle refers to the Ca9-treated cynomolgus monkey.

**Table. 1.** Summary of trio WGS and identification of DNMs.

| Monkey ID | Relationship | Treatment group | Depth | On-target ratio | Candidate DNMs | Final SNVs | Final INDELs |
| --- | --- | --- | --- | --- | --- | --- | --- |
| rmF<sup>wt</sup> | Father | wildtype | 45.81 | - |  | - | - |
| rmM1<sup>wt</sup> | Mother-1 | wildtype | 46.23 | - |  | - | - |
| rmM2<sup>wt</sup> | Mother-2 | wildtype | 45.23 | - |  | - | - |
| rmO1<sup>ko</sup> | Offspring-1 | cas9-treat | 48.39 | 0.435 | 7443 | 32 | 5 |
| rmO2-B<sup>ko</sup> | Offspring-2-brain | cas9-treat | 45.98 | 0.953 | 6603 | 30 | 3 |
| rmO2-M<sup>ko</sup> | Offspring-2-muscle | cas9-treat | 44.85 | 0.938 | 6698 | 28 | 2 |
| rmO2-L<sup>ko</sup> | Offspring-2-liver | cas9-treat | 45.35 | 0.952 | 6725 | 27 | 3 |
| rmO3<sup>ko</sup> | Offspring-3 | cas9-treat | 46.84 | 0.122 | 7100 | 34 | 1 |
| rmO4<sup>ko*</sup> | Offspring-4 | cas9-treat | 45.92 | 0 | 7209 | 26 | 1 |
| rmO5<sup>ko</sup> | Offspring-5 | cas9-treat | 47.60 | 0.321 | 7077 | 36 | 1 |
| cmF<sup>wt</sup> | Father | wildtype | 64.08 | - |  | - | - |
| cmM<sup>wt</sup> | Mother | wildtype | 76.11 | - |  | - | - |
| cmO<sup>ki</sup> | Offspring | cas9-treat | 68.42 | - | 11619 | 32 | 3 |

Using WGS data, we first reassessed the Cas9-editing efficiency at the *MCPH1* locus (Table 1, Supplementary Fig. 4). Consistent with the PCR-clone sequencing, we observed mosaic patterns of *MCPH1* knockout for all Cas9-edited monkeys. The knockout efficiency ranges from 12.2%-95.3% (Table 1) in the four Cas9-edited monkeys and no sequence change was detected in the knockout negative monkey (rmO4^*ko*\*^).

We then performed variant calling using four different tools, including GATK^21^, Samtools^22^, Freebayes^23^ and Platypus^24^ (Fig. 1b). Variant calling (VC), quality control (QC), site filtering (SF), genotype filtering (GF) and universal mask (UM) were performed to obtain high-confident variants (Methods) (Fig. 1b). The overlapped variants from the four different calling tools was taken as the high-confident variants (Supplementary Table 3)

With the use of the *SpeedSeq* pipeline^7^, we predicted the potential off-target locations for the two *MCPH1* sgRNAs. Among the 4,328 predicted off-target sites, no mutation was observed in the four Cas9-edited monkeys, including the monkey (rmO2^*ko*^) with multiple tissue samples (Supplementary Table 4). Hence, no off-target effects were detected at the predicted sgRNA binding sites.

To evaluate the genome-wide off-target effect, we explored the pattern of DNMs using the trio WGS data by TrioDeNovo software^25^ (Fig. 1b). We first validated the genetic relationship between the Cas9-treated monkeys and their parents^26^ (see Methods). The IBD (identity by descent) result agreed well with the original record (Supplementary Fig. 5). Initially, we obtained on average ~7,000 candidate DNMs for each Cas9-treated monkey by running TrioDeNovo (Table 1). We then performed multiple filtrations for each DNM set called by different tools (Methods). After overlapping the DNM sets, we obtained on average 32 high-confident DNMs for each monkey (30 substitutions and 2 indels) (Fig. 1c, 1d; Table 1; Supplementary Fig. 6,7; Supplementary Table 5). These high-confident DNMs can be explained by the spontaneous mutation rates of primates (0.98~2.17× 10^−8^ per nucleotide per generation) with 22~78 expected DNMs per generation^27, 28^. Consistently, we saw no correlation between the number of high-confident DNMs and the Cas9-editing efficiency (R^2^=-0.140, *P*=0.765, Pearson’s correlation test). In other words, the Cas9-editing efficiency does not affect the occurrence of DNMs in the Cas9-edited monkeys.

To further confirm the DNM pattern seen in rhesus monkeys, using the same pipeline, we analyzed the published trio WGS data of a gene-knockin model via CRISPR-Cas9 in cynomolgus monkeys^19^. The trio included an *Oct4-hrGFP* knockin cynomolgus monkey and his two wildtype parents (Fig. 1a, 1b). The results showed that only 35 DNMs (32 substitutions and 3 indels) were detected (Fig. 1c, 1d, Table 1), concordant with the pattern seen in rhesus monkeys. For the sgRNA-predicted off-target sites, only one mutation (a 2-bp deletion) was seen in the knockin cynomolgus monkey as reported in Cai et al.^19^.

In this study, we used four different tools to call variants and we took the overlap as the high-confident variant set. It is known that the performance of different tools varies when conducting genome-wide variant callings. Hence, a combination of different calling tools is necessary to identify high-confident variants from WGS data. Notably, the four different tools exhibited in general high consistency (>80%) for variant callings (Supplementary Table 3).

For DNMs identification, we initially identified ~7,000 candidate DNMs for each monkey. We found that >90% of them were the types disobeying the Mendel’s law due to unexpected allele combinations in the offspring monkeys, but the alleles were in fact present in the parents. For example, the genotypes of the parents are AA and GG respectively, and we saw AA or GG in the offspring instead of the expected genotype of AG. These candidate DNMs are most likely noise, not true DNMs, which may be caused by the technical bias of next-generation sequencing. The same scenario was also seen in the mouse data^13^. We filtered out these candidate DNMs by the DNM filtration procedure (“allele filtering”, Methods).

Finally, gene knockout and knockin are formed by different repair procedures in the cell. We analyzed trio sequencing data from both knockout (rhesus monkeys) and knockin (cynomolgus monkeys) monkey models, and we did not observe unexpected mutations in either model, suggesting that the homologous repair template does not induce off-target mutations. In conclusion, based on the WGS data from knockout and knockin monkeys, CRISPR-Cas9 can be considered a relatively safe gene editing system for primates.

## Methods

### Animals

All animals were housed at the AAALAC (Association for Assessment and Accreditation of Laboratory Animal Care) accredited facility of Primate Research Center of Kunming Institute of Zoology. All animal protocols were approved in advance by the Institutional Animal Care and Use Committee of Kunming Institute of Zoology (Approval No: SYDW-2010002).

### SgRNA design and *in vitro* transcription

Based on the rhesus monkey reference genome (Mmul_8.0.1), two sgRNAs were designed to target the *MCPH1* gene with sgRNA1 targeting exon2 and sgRNA2 targeting exon4. The sequences of the two sgRNAs are (PAM in bold): sgRNA1: CCTATGTTGAAGTGTGGTCATCC; sgRNA2: TTACACAGATGCAGGACAGCTGG. The sgRNAs were cloned into PUC57-sgRNA vector (Addgene No. 51132) (Supplementary Table 6). The sgRNAs were transcribed by MEGAshortscript Kit (Ambion, AM1354) after the vectors were linearized by DraI (NEB, R0129S). SgRNAs were purified by MEGAclear Kit (Ambion, AM1908). Cas9 mRNAs were transcribed by T7 Ultra Kit (Ambion, AM1345) after the pST1374-Cas9-NNLS-flag-linker vector (Addgene No. 44758) was linearized with AgeI (NEB, R0552S). Cas9 mRNAs were purified by RNeasy Mini Kit (Qiagen, 74104).

### Zygote injection and embryo transfer

Briefly, healthy female monkeys with regular menstrual cycles were used as oocyte donors for superovulation by intramuscular injection with rhFSH (Recombinant Human Follitropin Alfa, GONAL-F^®^, Merck Serono) for continuous 8 days, then rhCG (Recombinant Human Chorionic Gonadotropin Alfa, OVIDREL^®^, Merck Serono) on day 9. The oocytes were collected by laparoscopic follicular aspiration 36 hours after rhCG treatment. The MII (first polar body present) oocytes were selected for *in vitro* fertilization (IVF) and the fertilization was confirmed by the presence of two pronuclei. Fertilized eggs were injected with a mixture of Cas9 mRNA (20 ng/μl), sgRNA1 (10 ng/μl) and sgRNA2 (10 ng/μl) into cytoplasm using a Nikon microinjection system. The injected zygotes were cultured in the chemically defined, protein-free hamster embryo culture medium-9 (HECM-9) containing 10% fetal calf serum (Hyclone Laboratories, SH30088.02) at 37 °C in 5% CO2. The normally developed embryos from 2-cell to 8-cell with high quality were transferred into the oviduct of the matched recipients. A total of 6 monkeys were used as surrogate recipients, and typically, 4 embryos were transferred for each recipient female. The earliest pregnancy was diagnosed by ultrasonography about 30 days after transfer. Both pregnancy and number of fetuses were confirmed by fetal cardiac activity and presence of a yolk sac as detected by ultrasonography.

### DNA extraction

Genomic DNA was extracted using the DNeasy Blood & Tissue Kit (Qiagen, 69506) according to manufacturer’s instructions. The tissue samples included muscle, liver and brain from rmO1 and blood from the other monkeys. DNA samples were quantified using a NanoDrop spectrophotometer.

### Genotyping of *MCPH1* gene knockout rhesus monkeys

PCR primers were designed to amplify the sgRNA targeting region (Supplementary Table 7). Targeted fragments were amplified with Taq DNA polymerase from genomic DNA. PCR products were sub-cloned into pMD19 vector (Takara, 3271). The colonies were picked up randomly and sequenced by M13-F primer.

### Whole genome sequencing

Whole genome sequencing libraries were prepared using standard protocols for the Illumina X10 platform. Briefly, 100 ng DNA was fragmented using a Covaris LE220 (Covaris), size selected (300-550 bp), end-repaired, A-tailed, and adapter ligated. Libraries were sequenced using the Hiseq X Ten platform (Illumina) as paired-end 150 base reads. We generated on average 133 Gb raw sequence data per monkey. We performed quality control by fastqc and GATK, and the mean Q30 of read-pairs are higher than 88%. Each sample has a raw read depth (rDP) > 46. After masking the duplicates by Picard, we calculated the effective read depth (eDP) of the entire genome and the average eDP > 40. Mean percentage of PCR duplicates was lower than 13%, and the average mapped rate >99.1%. The properly paired reads are > 96% (Supplementary Table 2)

### Alignment and post-alignment processing

We used BWA MEM algorithm^29^ to perform alignment, where short reads of rhesus monkeys (*Macaca mulatta*) were mapped to their reference genome (genome build Mmul_8.0.1, rheMac8). The short reads of cynomolgus monkeys (*Macaca fascicularis*) were mapped to their reference genome (genome-build Macaca_fascicularis_5.0). The detailed command lines can be found in Supplementary Table 8. After the initial alignment, we run Picard’s MarkDuplicates to remove duplicates in both datasets.

### Variant Calling

We called single-nucleotide variants (SNVs), and insertions/deletions (INDELs), from de-duplicated bam files with GATK HaplotypeCaller^21^, Platypus^22^, Freebayes^23^ and Samtools^24^. The command lines can be found in Supplementary Table 8. For GATK, variants were called and a GVCF file was generated for each sample, and then joint calling were performed for GVCF files of each trio, separately. For Platypus, Freebayes and Samtools, we directly called the variants for each trio, separately.

### Variant filtering

The overview of the variant filtering strategies can be found in Fig. 1b. We used site filtering (SF), genotype filtering (GF) and universal masking (UM) to filter against variants with low quality in each VCF set called by different callers. SF strategy filters variants at the site level, which takes QD (variant confidence/quality by depth) (QD > 2.0), Mapping Quality (MQ > 30), Allele Bias (at least Pval < 0.05), Strand Bias (at least Pval < 0.05) into consideration. GF filters variants at the genotype level, which takes depth (15< DP <100 for SNVs) and genotype quality (GQ > 30) of each genotype into consideration. Since the information varies in the VCF sets generated by different variant calling tools, the corresponding SF and GF filtering variables and parameters are different. The details were summarized in Supplementary Table 9, 10. The UM is a sample independent mask that identifies complex regions in the human reference genome where variant calling can be challenging^30^. In our analysis, the UM included three components: (a) mappability mask; (b) low complexity regions; and (c) repeat regions. The command lines that generated (a) and (b) are provided in Supplementary Table 11. (c) were directly downloaded from UCSC. We merged the three sets of regions. SNVs and INDELs in the UM regions were filtered out, of which the detailed commands for can be found in Supplementary Table 9 and 10.

### Target region analysis

We extracted the reads, which aligned to the sgRNA binding regions as well as regions 100bp up-stream and down-stream by Samtools *tview^24^.* We then investigated the Cas9-target effect at these reads, and the number of reads with deletions near PAM were counted in calculating on-target rate (Supplementary Fig. 4).

### Prediction of off-target sites of *MCPH1* sgRNAs

As previous study described, genomic sites with “NGG” or “NAG” PAM motifs and ungapped alignment with up to 5 mismatches with sgRNA1 or sgRNA2 were treated as potential off-target sites^7^. We identified these potential off-target sites by in-house script and the results are listed in Supplementary Table 4.

### Relatedness validation

To reduce computational complexity and linkage disequilibrium (LD) effect, we only included the independent variants of whole-genome to create the IBS matrix by PLINK1.07^26^ with argument: *indep-pairwise 50 5 0.2*. A total of 866,199 independent variants were included in calculating IBD by PLINK1.07.

### DNMs calling

We adopted TrioDeNovo software to identified DNMs, using default settings and ran by each parent/offspring trio. After obtaining the results from running TrioDeNovo, we performed multiple filtrations to remove the false positive DNMs: 1) allele filtering: the DNM candidates where at least one allele was absent from parental genotypes; 2) dbSNP filtering: the DNM candidates must be absent from public SNV database (dbSNPBuildID=150); 3) cross filtering: the DNM candidates shared between offspring were removed. Finally, to decrease false positive rate, we took the overlapped set as the final DNMs detected in VCF sets of different callers (Fig. 1c, Supplementary Fig. 7).

### Data availability

The raw fastq data generated in this study was deposited to the Genome Sequence Achieve (http://gsa.big.ac.cn) and the accession number is CRA000954.

## Acknowledgements

We thank Yan Guo for her technical assistance in this study. We thank Xingxu Huang for providing plasmids. This study was supported by grants from the Strategic Priority Research Program (XDB13010000) and the Key Research Program of Frontier Sciences (QYZDJ-SSW-SYS009) of the Chinese Academy of Sciences, the National Natural Science Foundation of China (31730088, 91731303, 31525014, 31771388, and 31711530221), the Program of Shanghai Academic Research Leader (16XD1404700), the National Key Research and Development Program (2016YFC0906403), and the Shanghai Municipal Science and Technology Major Project (2017SHZDZX01).

## Author contributions

X.L. and B.S. designed the study; X.L., M.L., J.J., T.H., Y.H., X.M. performed experiments; X.H., L.Y., P.Z., X.Z. assisted in reproductive technique, microinjection and animal care; Y.H, C.Z., X.L., S.X. performed data analysis; W.J. provided the WGS data of the knockin monkeys; X.L., Y.H. and B.S. wrote the paper.

